# Hearing impairment is associated with enhanced neural tracking of the speech envelope

**DOI:** 10.1101/489237

**Authors:** Lien Decruy, Jonas Vanthornhout, Tom Francart

## Abstract

Elevated hearing thresholds in hearing impaired adults are usually compensated by providing amplification through a hearing aid. In spite of restoring hearing sensitivity, difficulties with understanding speech in noisy environments often remain. One main reason is that sensorineural hearing loss not only causes loss of audibility but also other deficits, including peripheral distortion but also central temporal processing deficits. To investigate the neural consequences of hearing impairment in the brain underlying speech-in-noise difficulties, we compared EEG responses to natural speech of 14 hearing impaired adults with those of 14 age-matched normal-hearing adults. We measured neural envelope tracking to sentences and a story masked by different levels of a stationary noise or competing talker. Despite their sensorineural hearing loss, hearing impaired adults showed higher neural envelope tracking of the target than the competing talker, similar to their normal-hearing peers. Furthermore, hearing impairment was related to an additional increase in neural envelope tracking of the target talker, suggesting that hearing impaired adults may have an enhanced sensitivity to envelope modulations or require a larger differential tracking of target versus competing talker to neurally segregate speech from noise. Lastly, both normal-hearing and hearing impaired participants showed an increase in neural envelope tracking with increasing speech understanding. Hence, our results open avenues towards new clinical applications, such as neuro-steered prostheses as well as objective and automatic measurements of speech understanding performance.

**Highlights:**

- Adults with hearing impairment can neurally segregate speech from background noise
- Hearing loss is related to enhanced neural envelope tracking of the target talker
- Neural envelope tracking has potential to objectively measure speech understanding

## 1. Introduction

Hearing impairment (HI) is one of the highest sources of disability and can be caused by many different etiologies (World Health Organization, 2018). The most common and well-known cause involves aging. With advancing age, a progressive deterioration occurs on different levels of the auditory system: from the sensors in the cochlea, stria vascularis, spiral ganglion neurons up to the neural pathways of the central auditory system, leading to a sensorineural hearing loss (Bess and Humes, 1995; Schmiedt, 2010). Depending on which of these declines occurs or dominates, different audiometric phenotypes of age-related hearing loss can be distinguished (Dubno et al., 2013). Next to age-related hearing loss, sensorineural hearing loss can also occur in young adults due to prenatal or postnatal diseases (e.g. cytomegalovirus), genetic factors and occupational or recreational noise exposure (Bess and Humes, 1995; World Health Organization, 2018).

Sensorineural hearing loss is mainly characterized by a decreased hearing sensitivity of the incoming sound, but it also leads to other deficits that often cannot be overcome by hearing aids (Dillon, 2001; Humes et al., 2013). At the peripheral level, outer hair cell dysfunction also causes reduced frequency resolution, resulting in a distorted output of the cochlea which makes a segregation between speech and noise at the subsequent levels of the auditory system more difficult (Dillon, 2001; Hopkins and Moore, 2011). In addition, several studies have shown that HI adults benefit less from listening in the gaps when speech is embedded in the presence of fluctuating maskers (Festen and Plomp, 1990; Goossens et al., 2017; Shinn-Cunningham and Best, 2008). It is suggested that a deficient temporal processing of both low- (envelope) and high-frequency (temporal fine-structure; TFS) modulations underlie these speech-in-noise difficulties. Behavioral studies have demonstrated that HI adults obtain lower, better thresholds when performing an amplitude modulation detection task, indicating an enhanced envelope sensitivity when having a HI (Füllgrabe et al., 2003; Wallaert et al., 2017). In contrast to the envelope, HI adults have shown a decreased sensitivity to TFS (Hopkins and Moore, 2011). Accordingly, animal studies revealed that auditory nerve fibers show enhanced envelope (Henry et al., 2014; Kale and Heinz, 2010) but degraded TFS encoding (Henry and Heinz, 2012) in mammals with noise-induced hearing loss. Although the peripheral effects of hearing impairment on speech perception have been extensively investigated, a lot remains unknown about the consequences of sensorineural hearing loss at the higher levels of the auditory system.

A few electrophysiological studies have assessed this by recording auditory evoked potentials (AEPs) to non-speech or short speech stimuli. Both at the brainstem and cortical level, enhanced AEPs have been observed for HI adults. For example, Anderson et al. (2013) recorded frequency-following responses (FFR) to a /da/ stimulus and found enhanced envelope encoding for HI older adults compared to NH peers. Goossens et al. (2019) found significantly higher brainstem as well as cortical auditory steady-state responses (ASSRs) for young (20-30 years) and middle-aged (50-60 years), but not for older HI adults (70-80 years old). Similarly, sensorineural hearing loss is demonstrated to relate with enhanced magnetoencephalography (MEG) responses for amplitude modulated noise in primary auditory regions (Millman et al., 2017) and increased cortical AEPs (CAEPs) originating from frontal regions (Campbell and Sharma, 2013). Although these studies provide evidence for enhanced envelope sensitivity in HI adults, research also showed decreased subcortical responses or no significant differences in cortical CAEPs between NH and HI adults (Ananthakrishnan et al., 2016; Billings et al., 2015; Koerner and Zhang, 2018).

A better understanding of the neural changes underlying speech-in-noise difficulties in HI adults, could be achieved by measuring neural activity to natural, continuous speech instead of analyzing responses to short, artificial non-speech sounds or syllables. Over the past decade, a growing body of research has shown that when persons listen to stories or sentences, their neural activity tracks the speech envelope of the acoustic stimulus (Ding and Simon, 2014; Lalor and Foxe, 2010). Moreover, studies investigating the cocktail party problem, i.e. ability to attend to one talker while ignoring others, have shown an increased neural tracking of the speech envelope for the target versus competing talker in NH listeners (Das et al., 2016; Ding and Simon, 2012; O’Sullivan et al., 2015). To our knowledge, only three studies have been published which assessed neural tracking of the speech envelope in HI listeners. Petersen et al. (2017) measured neural tracking of the speech envelope in older HI adults with a varying degree of hearing loss, using electroencephalography (EEG). They showed that their neural activity tracks the target talker better than the competing talker similarly to adults without hearing loss. Nonetheless, worse hearing loss was associated with higher neural tracking of the speech envelope for the ignored, competing talker, resulting in a weaker differential tracking for target versus competing talker for HI adults. Although Presacco et al. (2019) and Mirkovic et al. (2019) also found a similar robust neural tracking of the target versus competing talker speech envelope for HI adults, no significant differences in envelope tracking were reported between NH and HI adults in these studies. To improve further readability of the article, we will use the term “(neural) envelope tracking” throughout the next sections, when referring to neural tracking of the speech envelope.

Because inconsistent findings have been shown about the effect of hearing impairment on brain responses to non-speech stimuli (Ananthakrishnan et al., 2016; Billings et al., 2015; Campbell and Sharma, 2013; Goossens et al., 2019; Koerner and Zhang, 2018; Millman et al., 2017) and natural speech (Mirkovic et al., 2019; Petersen et al., 2017; Presacco et al., 2019), the first aim of the present study was to assess the neural consequences of hearing impairment. We did this by comparing neural envelope tracking of a target and competing talker for 14 HI participants with data of 14 NH adults collected in a previous study (Decruy et al., 2019). Previous studies have shown that neural envelope tracking increases with advancing age in NH adults (Decruy et al., 2019; Presacco et al., 2016). Since sensorineural hearing loss becomes more prevalent with advancing age, it is thus important to carefully age-match the NH and HI participants to assess the specific effect of hearing impairment on neural envelope tracking, without the confound of age. Next to neural envelope tracking of natural speech, we also measured cortical ASSRs to tone pips and hypothesize based on the findings of Millman et al. (2017) and Goossens et al. (2019) that HI adults show enhanced cortical responses relative to their NH peers.

Finally, recent studies have shown the potential of measures of neural envelope tracking to predict speech understanding performance (Lesenfants et al., 2019; Vanthornhout et al., 2018). These studies, however, were performed in NH participants. Closely related to this, two recent studies have investigated neural envelope tracking as a function of SNR in HI adults but found contradicting results. Petersen et al. (2017) found that neural envelope tracking did not increase with changes in SNR for persons with a higher degree of hearing impairment whereas Presacco et al. (2019) found an increase in envelope tracking with increasing SNR for HI adults, but not for NH adults. A plausible reason for these inconsistent results could be the choice of SNR when testing NH and HI adults. More specifically, HI adults have worse speech understanding than NH adults and presenting the same SNRs to these two populations could lead to ceiling or floor effects for one of the populations. To overcome this, the second aim of this study was to examine the relation between speech understanding and neural envelope tracking using more direct speech understanding measures (percentage correctly recalled sentences or ratings), in addition to presenting subject-specific SNRs. Using this approach, we could evaluate whether HI adults also show an increase in neural envelope tracking as a function of speech understanding, despite having a hearing loss.

## 2. Material and methods

### 2.1 Participants

Fourteen adults (8 female) with sensorineural hearing loss, aged between 21 and 80 years old, participated in the present study. To investigate the effect of hearing impairment on neural envelope tracking, the results of these participants were compared to those of fourteen age-matched NH adults collected during a previous study (age-matching table 1, Decruy et al., 2019). All participants had Dutch (Flemish) as their mother tongue, except one HI person (P1) who grew up in the Netherlands but had resided in Flanders the past four years. To ensure that decreased hearing sensitivity in the HI adults was due to a sensorineural and not conductive hearing loss, both air and bone conduction thresholds were obtained. The individual hearing thresholds demonstrate that hearing loss was most apparent in the high frequencies and varied among HI participants from mild to severe hearing loss (figure 1). From the tonal audiograms, it can also be inferred that both NH and HI participants had symmetrical hearing. Symmetry was verified based on the criteria derived from the AMCLASS algorithm of Margolis and Saly (2008). As can be inferred from table 1, all participants wore bilateral hearing aids to compensate for their hearing loss.

**Figure 1:**
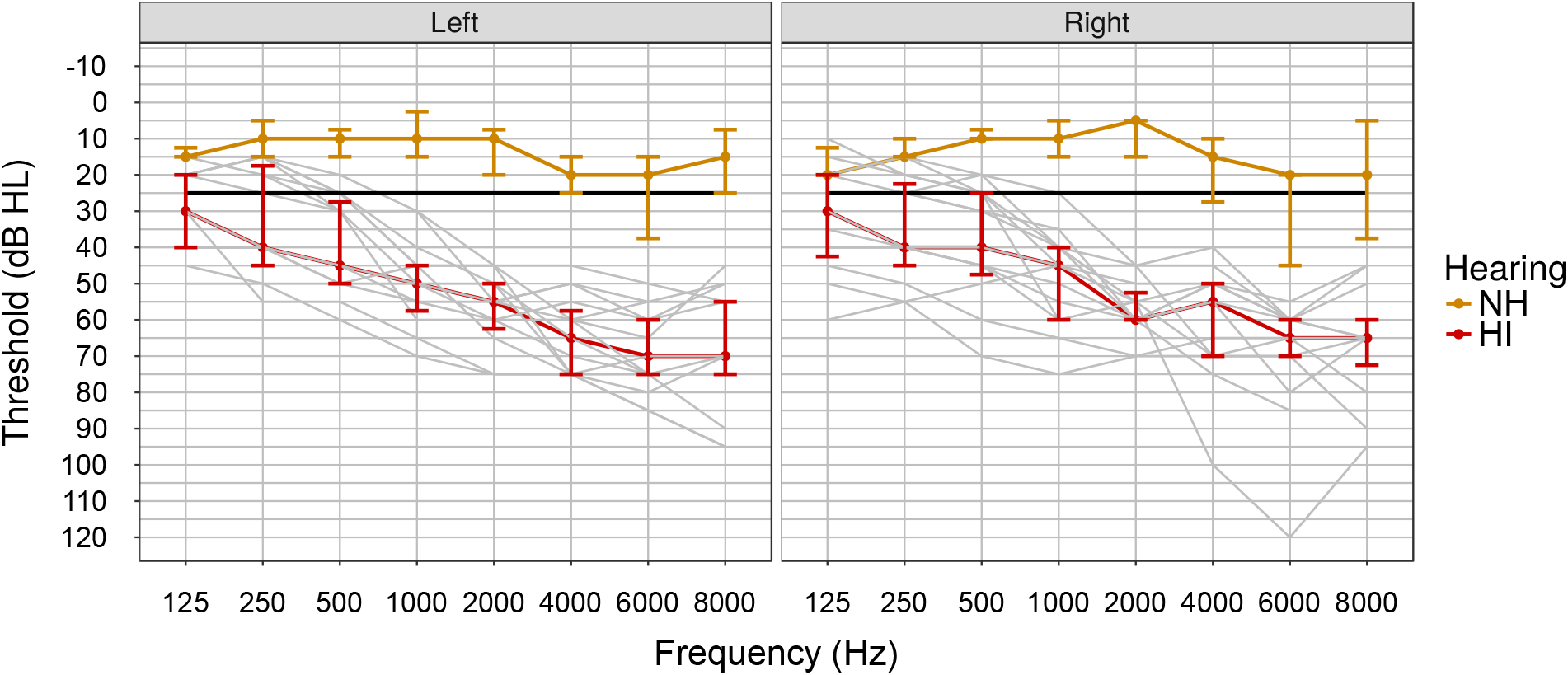
Median air conduction thresholds (in dB HL) of normal-hearing and hearing impaired participants. Error bars indicate the interquartile range. Individual audiometric curves for hearing impaired participants are also depicted.

**Table 1:**
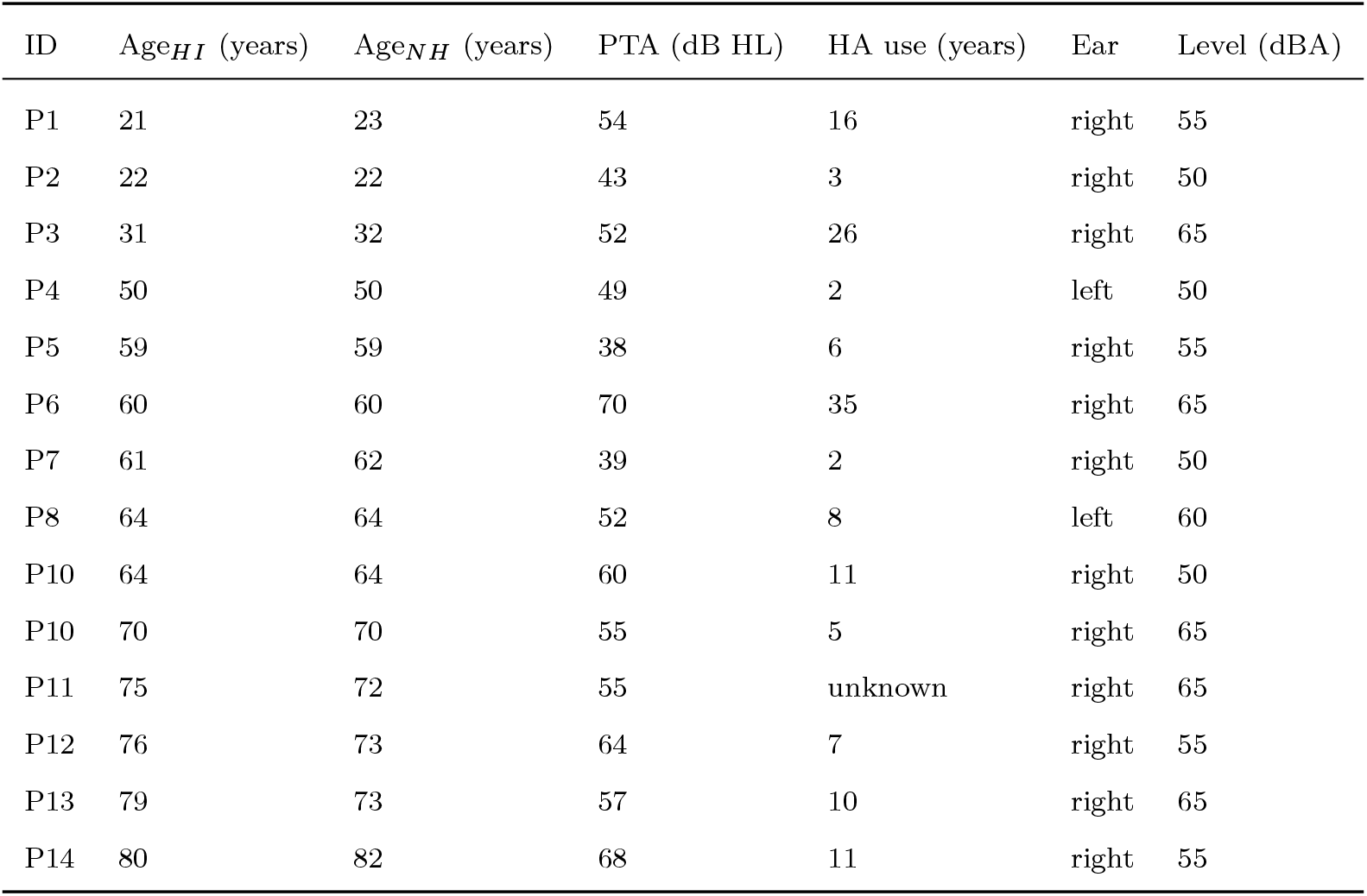
Characteristics of hearing impaired (HI) participants and age-matched normal-hearing (NH) adults. Per HI participant, the age, age of matched NH adult, pure tone average (PTA) of four frequencies (500, 1000, 2000, 4000 Hz), hearing aid (HA) use, stimulation side (ear) and subject-specific stimulation intensity (level) for speech stimuli after providing linear amplification (see 2.2.4), are reported. One HI participant had worn hearing aids for several years but could not remember the precise number of years.

| ID | Age<sub>HI</sub> (years) | Age<sub>NH</sub> (years) | PTA (dB HL) | HA use (years) | Ear | Level (dBA) |
| --- | --- | --- | --- | --- | --- | --- |
| P1 | 21 | 23 | 54 | 16 | right | 55 |
| P2 | 22 | 22 | 43 | 3 | right | 50 |
| P3 | 31 | 32 | 52 | 26 | right | 65 |
| P4 | 50 | 50 | 49 | 2 | left | 50 |
| P5 | 59 | 59 | 38 | 6 | right | 55 |
| P6 | 60 | 60 | 70 | 35 | right | 65 |
| P7 | 61 | 62 | 39 | 2 | right | 50 |
| P8 | 64 | 64 | 52 | 8 | left | 60 |
| P10 | 64 | 64 | 60 | 11 | right | 50 |
| P10 | 70 | 70 | 55 | 5 | right | 65 |
| P11 | 75 | 72 | 55 | unknown | right | 65 |
| P12 | 76 | 73 | 64 | 7 | right | 55 |
| P13 | 79 | 73 | 57 | 10 | right | 65 |
| P14 | 80 | 82 | 68 | 11 | right | 55 |

In addition to pure tone audiometry, an interview was administered to question medical history, education, visual acuity, onset, etiology and duration of hearing loss, hearing aid use and laterality preference (table 1; Coren, 1993). None of the participants reported a medical condition, such as serious concussions or taking sleeping medication (Van Lier et al., 2004), that could bias our results. Participants also did not report problems with color vision or visual acuity that could not be corrected by wearing glasses or contact lenses. Furthermore, no diagnoses of learning disorders were reported. We checked this because research has shown that dyslexia can affect auditory brain responses (De Vos et al., 2017; Poelmans et al., 2012). Lastly, all HI adults passed cognitive screening as they obtained a score equal or higher than 26/30 on the Montreal Cognitive Assessment (Dautzenberg and Jonghe, 2004; Nasreddine et al., 2005). The study was approved by the Medical Ethics Committee UZ KU Leuven / Research (reference no. S57102 (Belg. Regnr: B322201422186)). HI participants gave their written consent and were paid for participation.

### 2.2 Behavioral experiments

In the present study, HI participants completed two similar main experiments as the NH adults (Decruy et al., 2019). As shown in overview figure 2, three behavioral experiments were administered in a first session of approximately two and a half hours. After pure tone audiometry, linear frequency-specific amplification was determined per HI participant. Then, speech understanding was evaluated using the Flemish matrix sentence test. Lastly, cognitive tests were administered to assess the interplay of cognition, hearing impairment and neural envelope tracking. Below, the different behavioral experiments are briefly described. For more details, we refer to Decruy et al. (2019).

**Figure 2:**
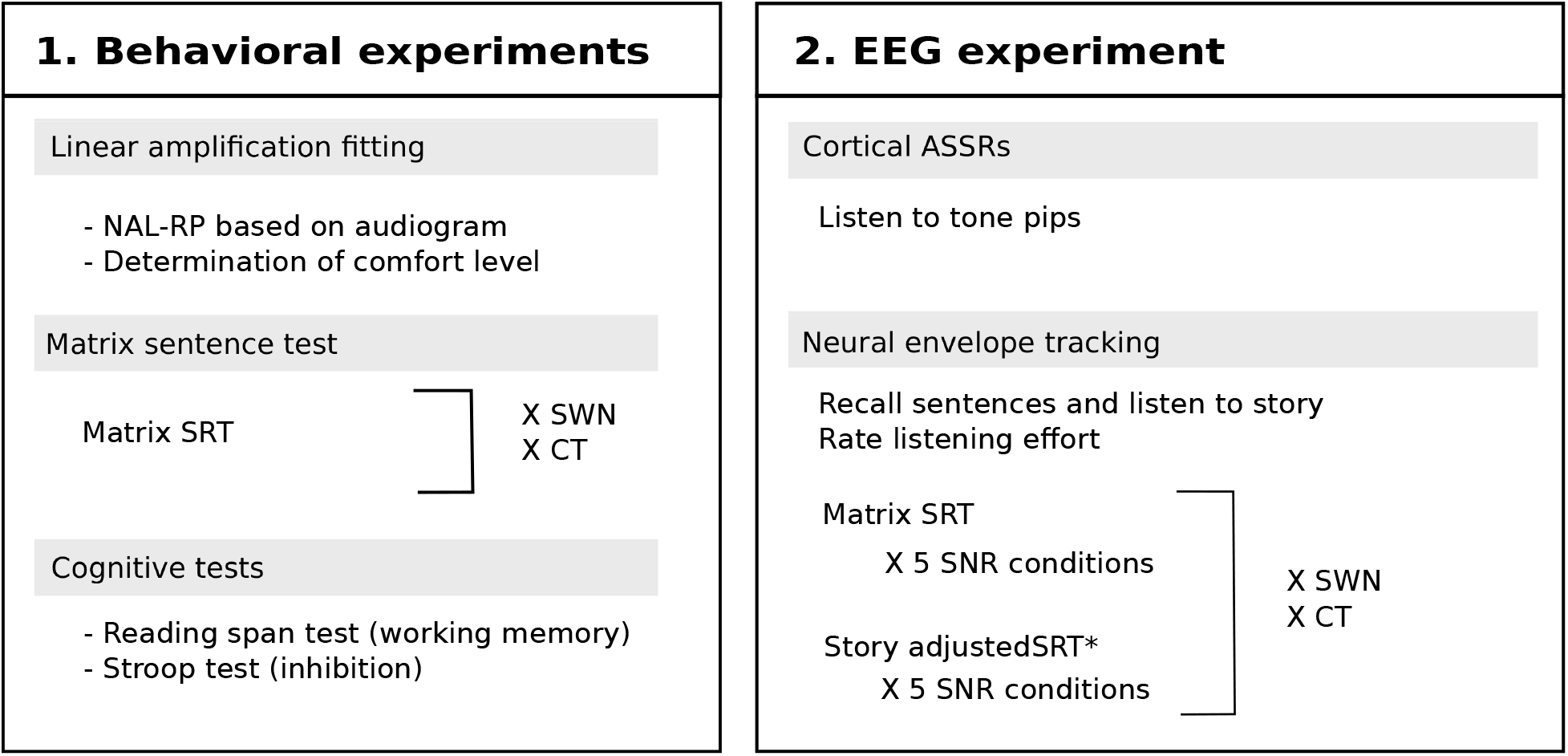
Overview of the main procedures. The levels for the story were created based on an adjusted version of the self-assessed Békesy procedure (Decruy et al., 2018; Decruy et al., 2019)

#### 2.2.1 Linear amplification

To investigate the effects of hearing impairment regardless of audibility, we provided our HI participants with subject-specific, linear frequency-specific amplification according to National Acoustics Laboratory-Revised Profound algorithm (NAL-RP) (Byrne et al., 2001, 1990; Dillon, 2001). We preferred linear amplification because compression can alter the envelope of speech, making it difficult to investigate the specific effects of hearing impairment on neural envelope tracking. Frequency-specific amplification was implemented by filtering the stimuli by a 512-coefficient finite impulse response filter which was determined in MATLAB R2016b based on the individual hearing thresholds.

#### 2.2.2 Speech understanding in noise

During the Flemish matrix sentence test, participants were instructed to recall five-word sentences spoken by a female talker (e.g. “Sofie ziet drie blauwe dozen” (“Sofie sees three blue boxes”)). These sentences were embedded in a stationary speech weighted noise (SWN) or a story narrated by a male talker (competing talker (CT)). Using the adaptive procedure of Brand and Kollmeier (2002), the level of the masker was adjusted based on the participant’s response to converge to 50% speech understanding, also called the speech reception threshold (SRT; Luts et al., 2014). The SRT was defined as the last SNR presented in a list of 20 sentences. Per type of masker (SWN and CT), three lists were conducted. To avoid confounds of procedural learning, only the SRT of the third list was used as the matrix SRT in our analyses (Luts et al., 2014).

Next to evaluating speech understanding performance, the matrix SRTs were used to determine subject-specific SNRs for the EEG experiment. In contrast to Petersen et al. (2017), we did not present the story at the same SNRs as the matrix sentences because we recently demonstrated significant differences between the SRTs for standardized sentences and stories (Decruy et al., 2018). Using an adapted version of the self-assessed Békesy procedure, we determined the adjustment values needed to create the SNRs for the story during the EEG-experiment (story adjusted SRT; figure 2; Decruy et al., 2019).

#### 2.2.3 Cognitive tests

There has been keen interest in the relation between hearing impairment and cognition because it is been suggested that persons with decreased sensory processing have an increased risk of developing cognitive impairment, such as dementia (Lin et al., 2013; Rönnberg et al., 2016). In addition, research has also shown an association between worse cognitive performance and disabling hearing loss in older adults without indication of cognitive impairment (Humes et al., 2013; Lin et al., 2011). To investigate the interplay between hearing impairment, neural envelope tracking and cognition, our HI participants completed both a working memory and inhibition test since studies have suggested that these cognitive skills could underlie the speech-in-noise difficulties related to hearing impairment (Akeroyd, 2008; Petersen et al., 2017; Shinn-Cunningham and Best, 2008). The Flemish computerized version of the Reading Span Test (RST; Van Den Noort et al., 2008; Vercammen et al., 2017) assesses working memory by measuring a person’s ability to remember sentence-final words of subsets of 2, 3, 4, 5 or 6 sentences that they had to read out loud. The Stroop Test (Hammes, 1978) evaluates inhibition skills by comparing the results on a congruent task in which persons have to name the colors of rectangles, and a incongruent task in which the color of words, e.g. “blue” printed in red, have to be named (red) while inhibiting reading the word (blue). More details about these cognitive tests are described by Decruy et al. (2019).

#### 2.2.4 Apparatus and presentation of stimuli

For all participants, speech stimuli were presented monaurally through ER-3A insert phones (Etymotic Research, Inc., IL, USA) using the software platform APEX (Dept. Neurosciences, KU Leuven) (table 1; Francart et al., 2008). The masker story was set to the same root mean square level and spectrum as the matrix sentences, and silences were shortened to a maximum duration of 200 ms (Decruy et al., 2019). For NH participants, the level of the target speech stimuli was fixed at an intensity of 55 dB SPL (A weighted). For HI adults, on the other hand, speech stimuli were presented at a subject-specific level. The NAL-RP algorithm estimates the needed linear amplification based on the individual hearing thresholds, but does not guarantee a comfortable level. This level was determined by varying the overall intensity of a story in quiet until participants indicated on a scale that it was minimally effortful and comfortable and reported that they could understand approximately 100% of the story (table 1). This is similar to a volume control button, a common feature of a hearing aid. The level of the masker was adjusted according to the target signal-to-noise ratio (SNR). All stimuli were calibrated with a type 2260 sound level pressure meter, a type 4189 half-inch microphone and a 2cc coupler (Bruël & Kjaer, Copenhagen, Denmark).

### 2.3 EEG experiment

In a second session of approximately three hours with breaks, we started the EEG experiment by recording ASSRs to tone pips (overview figure 2). Next, for both SWN and CT, EEG was measured while HI participants listened to matrix sentences or a story presented at five different SNRs to estimate neural envelope tracking as a function of speech understanding. The matrix sentences and story were both spoken by a female speaker. A story was also presented because recalling and correctly identifying matrix sentences does not reflect all processes involved in understanding natural speech. Listening to a story and answering questions about the content, on the other hand, activates more high-level cognitive processes such as using context cues to disentangle the target speech from the masker. We chose to not further address this difference in this manuscript and will use “speech understanding” to refer to both recalling of matrix sentences as well as listening to the story. Lastly, the 12 minutes story “Milan”, narrated by a male speaker, was presented without masker to obtain training data for the linear decoder (see 2.3.2).

#### 2.3.1 Procedure

Similar to Petersen et al. (2017), we created four subject-specific SNRs that covered the psychometric function of each individual by raising and lowering the subject-specific matrix and story adjusted SRT (50% SU) obtained during the behavioral experiment. For SWN, we used a step size of 3 dB to create the four subject-specific SNRs as follows: 20% (SRT - 3 dB), 50% (SRT), 80% (SRT + 3 dB) and 95% SU (SRT + 6 dB). For CT, we created the same four SU levels, but used a step size of 4 dB because the use of stories as a competing masker has been shown to result in less steep psychometric functions compared to stationary SWN maskers (Francart et al., 2011; MacPherson and Akeroyd, 2014). In addition to these four subject-specific SNRs, the target speech was also presented at two fixed SNRs conditions: without masker (No noise) and at the same level of the competing talker (0 dB SNR). In total, HI adults thus completed 20 conditions (2 maskers x 2 speech materials x 5 SNRs) of which the order of the four “speech material x masker” blocks (e.g. matrixCT, storySWN, matrixSWN, storyCT) as well as the five SNR conditions within each block were randomized across participants. For the matrix sentences, one list of 20 sentences was presented at the given subject-specific or fixed SNR. For the story, one part of approximately 3 minutes of “De Wilde Zwanen” from Hans Christian Andersen, was presented continuously at the subject-specific or fixed SNR.

To investigate the relation between speech understanding (SU) and envelope tracking, HI participants were asked to recall each matrix sentence out loud. Since a story is continuous, recalling the full segment was not feasible and therefore we asked participants to rate their speech understanding after each SNR condition using a scale from 0 to 100%. This rating was also performed for the matrix sentences in order to correct for subjective bias. More specifically, we adjusted the story ratings by adding the difference score between the percentage of correctly *recalled* matrix sentences and the *rated* percentage for the matrix sentences. Using the same scale, self-reported listening effort was also assessed after each matrix or story condition by asking the participants “How much effort do you need to expend to understand the sentences/story?”, with 100% indicating “extreme effort” and 0% indicating “no effort”.

In addition to the speech stimuli, we also measured cortical ASSRs to tone pips to assess the effect of hearing impairment on non-speech sounds. The tone pips were created in MATLAB R2016b, using a 21 ms sinusoid (with 4 ms on- and off ramps) with a carrier frequency of 500 Hz. The tone pip was repeated 250 times to obtain a repetition frequency of approximately 1.92 Hz. In contrast to the NH participants, the tone pips in this study were not presented at 90 dBpeSPL, but were linearly amplified per participant according to NAL-RP (see 2.2.1).

#### 2.3.2 Signal processing

In the next paragraphs, the several signal processing steps to measure neural envelope tracking and ASSRs to tone pips are described (for details see Decruy et al., 2019; Vanthornhout et al., 2018).

With regard to neural envelope tracking, we did not use the crosscorrelation approach of Petersen et al. (2017) in which a cluster of EEG electrodes were selected based on statistics. Instead, we used a backward decoding model which is a data-driven approach that uses a linear spatiotemporal filter or decoder that optimally combines the different EEG signals in order to reconstruct a single channel speech envelope (Crosse et al., 2016; Lalor et al., 2009). The several steps to obtain our measure of neural envelope tracking using this method, are the following. First, we extracted the envelopes from the original, unamplified, speech stimuli. Then, we filtered the envelopes and EEG from 1 to 8 Hz (see Decruy et al., 2019). Second, we trained a linear decoder that combines the signals of 64 EEG channels and their time shifted versions (integration window 0-500 ms), on the EEG responses to the story “Milan” presented without masker (mTRF toolbox: Lalor et al., 2006, 2009). The decoder was then applied on the EEG responses to the matrix sentences and story in the five SNR conditions to obtain a reconstructed envelope. Neural envelope tracking of the target and competing talker was measured by correlating the reconstructed envelope with the actual target and competing envelope. Lastly, we calculated a significance level of the correlation by correlating random permutations of the actual and reconstructed envelope and taking percentile 2.5 and 97.5 to obtain a 95% confidence interval.

Since HI participants listened to linearly amplified speech stimuli, we checked whether neural envelope tracking changed when calculating the correlation between the reconstructed envelope and the amplified versus unamplified envelope of the target or competing stimulus. According to our expectations, no significant difference in correlation was detected using two Linear Mixed-effect Models with participant as random effect, neural envelope tracking measure for target (model 1) or competing talker (model 2) as outcome measure and type of envelope (amplified versus unamplified) as fixed-effect term (model 1 / target talker: *β* = 1.18e-04, SE = 5.05e-03, p = 0.981; model 2 / competing talker: *β* = −7.69e-06, SE = 5.82e-03, p = 0.999). Based on these results, we decided to only report the results for the unamplified envelopes.

To analyze the ASSRs evoked by the tone pips, we applied denoising source separation (DSS) on high-pass filtered EEG-data (cut-off frequency: 0.5 Hz). DSS is a data-driven algorithm based on principal component analysis that designs a spatial filter that separates neural activity into stimulus-related and stimulus-unrelated components, based on a criterion of stimulus-evoked reproducibility (Cheveigné and Simon, 2008). This way, no subset of electrodes had to be selected. Then, we segmented the EEG-signals in epochs of approximately 2 s, removed artifacts using a fixed percentile criterion of 95% and transformed the epochs into the frequency domain using a Fast Fourier Transformation. The size of the ASSR was calculated as follows: 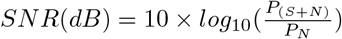 with *P_N_* reflecting the power of the non-synchronized neural activity and *P*_(*S*+*N*)_ reflecting the total power of the synchronized neural response to the tone pip and EEG noise in the frequency bin of interest (1.92 Hz).

#### 2.3.3 EEG recording

For all HI participants, EEG was recorded in a triple-walled, soundproof booth with Faraday cage at ExpORL. The same apparatus and presentation set-up as the behavioral experiment was used. More specifically, participants listened to the matrix sentences, story and tone pips monaurally through ER-3A insert phones. The stories, target and masker, were set to the same root mean square level and spectrum as the matrix sentences, and silences were shortened to a maximum duration of 200 ms (Decruy et al., 2019). EEG-data was collected using a BioSemi ActiveTwo recording system (Amsterdam, Netherlands) with 64 active Ag/AgCl electrodes and two extra electrodes, serving as the common electrode (CMS) and current return path (DRL). The 66 electrodes were mounted in head caps, designed according to a 10-20 system. Using the BioSemi ActiView software, EEG was continuously recorded, digitized at a sampling rate of 8192 Hz and stored on a hard disk for off-line signal analysis in MATLAB R2016b.

### 2.4 Statistical analysis

Using R software (version 3.4.4; nlme package - version 3.1-131.1; Field et al., 2012; Pinheiro et al., 2017), we investigated the effect of hearing impairment on neural envelope tracking by means of Linear Mixed-effect Models (LMMs) in which all the collected data points were included. The fixed effects in the LMMs consisted of the predictors of interest whereas the random effects included the variable participant, nested in one of the repeated measures predictors if this improved the model fit. Linear Fixed-effect Models (LFMs) were used to investigate the effects of hearing loss on cortical ASSRs to tone pips and cognition. All models were fitted using the maximum likelihood estimation. The best fitting model was determined by progressively introducing multiple fixed effects and the corresponding interactions and comparing the different models using likelihood ratio tests and Akaike’s Information Criterion (AIC; Akaike, 1974). The outcomes of the final model are discussed in the results by reporting the unstandardized regression coefficient (*β*) with standard error (SE), degrees of freedom (df), t-Ratio and p-value per fixed-effect term. For all models, we used a significance level of *α* = 0.05 unless we used post hoc tests to further investigate interaction effects (e.g. correction for multiple comparisons using the method of Holm, 1979). For one NH participant, we did not obtain neural envelope tracking at one condition (0 dB SNR). For the analysis of this SNR condition, we also excluded the data of the age-matched HI participant (P1).

## 3. Results

### 3.1 Speech understanding in noise

We first assessed the effect of hearing impairment on speech understanding in noise by comparing the matrix SRTs between NH and HI adults for two masker types, SWN and CT (LMM with three predictors: hearing, age, masker and their interactions, see table 2; figure 3). Post hoc tests revealed a more detrimental effect of hearing impairment when speech was embedded in CT (mean difference between NH CT vs HI CT = −11.1 dB, p < 0.001) compared to SWN (mean difference between NH SWN vs HI SWN = −2.8 dB, p = 0.048). In addition, we also found a significant increase in SRT with advancing age that was steeper for CT versus SWN (p = 0.001; table 2).

**Table 2:** Linear Mixed-effect Model: The effect of hearing impairment (hearing), age and masker type on the matrix speech reception threshold. Regression coefficients (*β* values), standard errors (SE), degrees of freedom (df), t-Ratios and p-values are reported per fixed-effect term. Participant was included as a random effect. Interactions are indicated by the symbol “:”.

| Fixed-effect terms | $\beta$ value | SE | df | t-Ratio | p-value |
| --- | --- | --- | --- | --- | --- |
| Intercept(for NH/SWN) | -11.76 | 1.97 | 25 | -5.97 | < 0.001 |
| Hearing | 2.80 | 1.17 | 25 | 2.40 | 0.024 |
| Age | 0.07 | 0.03 | 25 | 2.41 | 0.024 |
| Masker | -18.53 | 2.29 | 25 | -8.09 | < 0.001 |
| Hearing:Masker | 8.26 | 1.36 | 25 | 6.09 | < 0.001 |
| Age:Masker | 0.13 | 0.04 | 25 | 3.73 | 0.001 |

**Figure 3:**
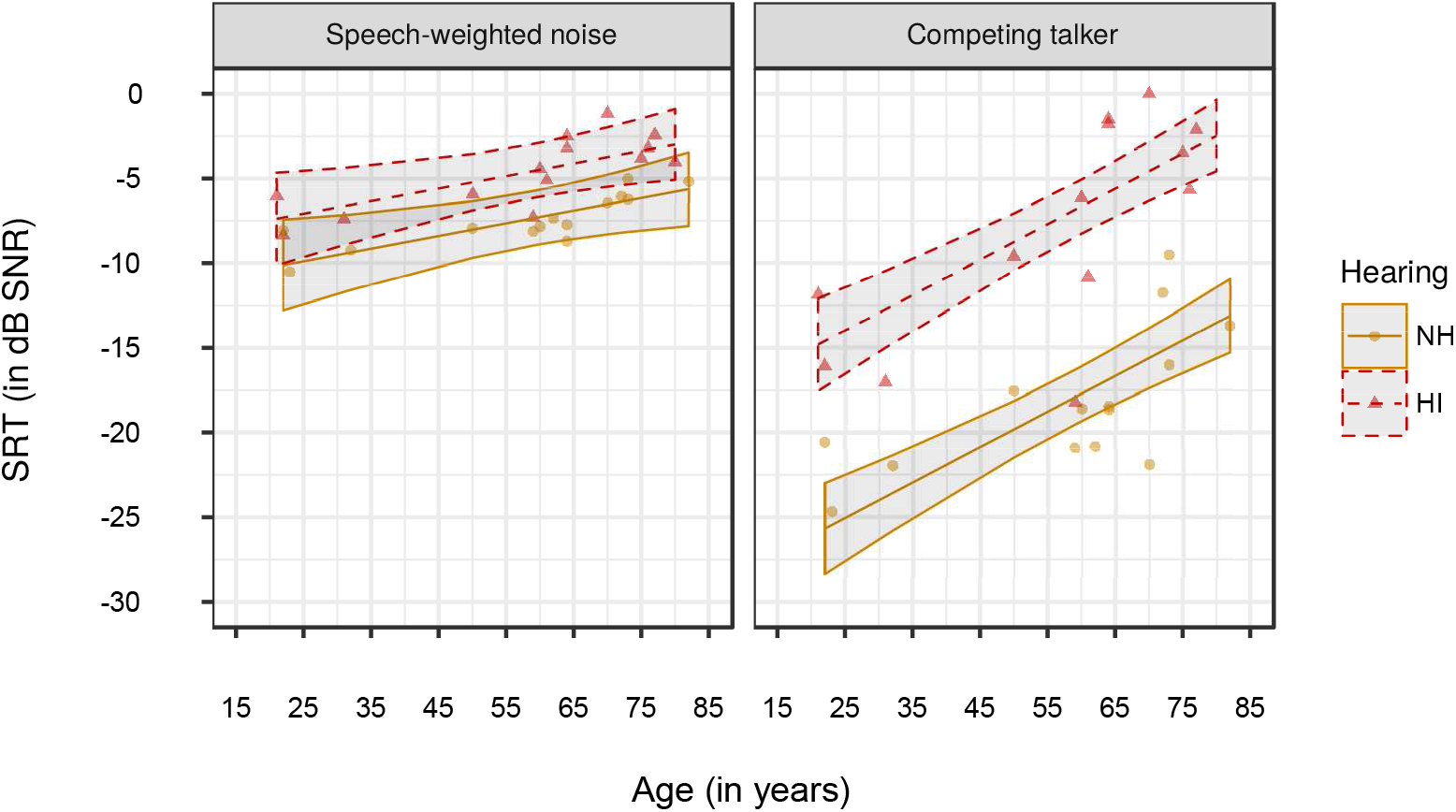
The matrix speech reception threshold (SRT) for 14 HI and 14 age-matched NH adults, in two masker types: speech-weighted noise (SWN) and a competing talker (CT). Per masker type, regression lines with confidence intervals (shaded areas) were fitted on all the data points for NH and HI adults (color-coded).

### 3.2 Link between speech understanding and neural envelope tracking

For both NH and HI adults, neural envelope tracking was measured as a function of speech understanding (SU). An LMM with six predictors (SU, hearing, age, age^2^, speech material and masker) and their interactions was used to predict neural envelope tracking for NH and HI adults. As shown in figure 4, we found that neural envelope tracking increased with increasing speech understanding for both NH and HI participants (p = 0.001; table 3). In addition, the LMM revealed two significant interaction effects with speech understanding: envelope tracking increased more steeply (1) with increasing speech understanding for older adults (p = 0.028; table 3) and (2) for matrix sentences compared to the story (p = 0.003; table 3).

**Figure 4:**
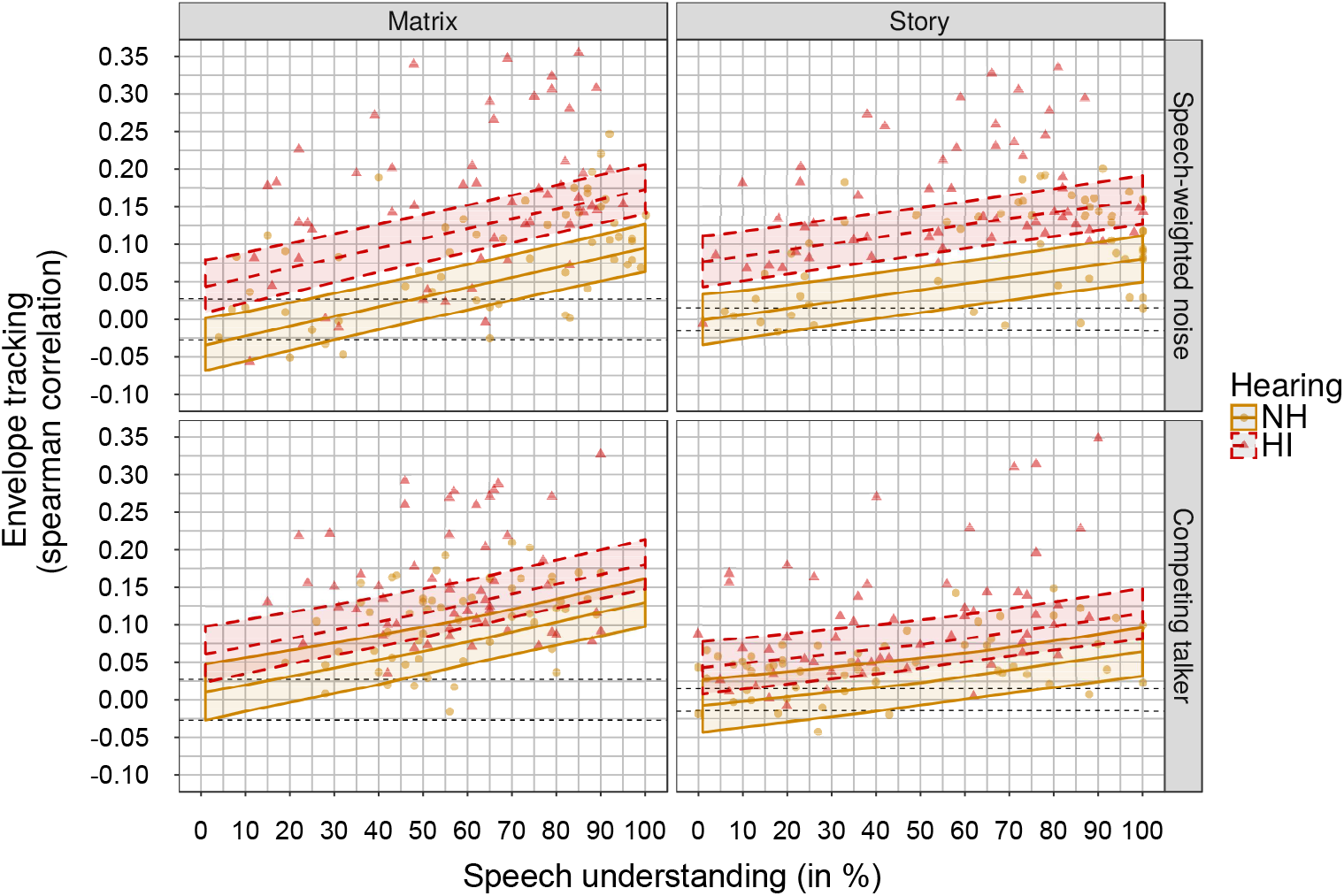
Neural tracking of the envelope as a function of speech understanding, measured in 14 HI and 14 age-matched NH adults. Per type of speech material and masker, two regression lines with confidence intervals (shaded areas) were fitted on all subject-specific data points for NH and HI participants (color-coded). The dashed black lines indicate the significance level.

**Table 3:** Linear Mixed-effect Model: The effect of speech understanding (SU; subject-specific SNRs), hearing impairment (hearing), age, type of speech material and masker on neural envelope tracking. Regression coefficients (*β* values), standard errors (SE), degrees of freedom (df), t-Ratios and p-values are reported per fixed-effect term. The variable participant nested in SU-level, was included as a random effect. Interactions are indicated by the symbol “:”.

| Fixed-effect terms | $\beta$ value | SE | df | t-Ratio | p-value |
| --- | --- | --- | --- | --- | --- |
| Intercept(for NH/Matrix/SWN) | 0.04 | 0.06 | 328 | 0.68 | 0.494 |
| SU | 8.55e-04 | 2.59e-04 | 328 | 3.30 | 0.001 |
| Speech material | 0.03 | 0.01 | 328 | 3.01 | 0.003 |
| Age | -4.49e-03 | 2.55e-03 | 24 | -1.76 | 0.091 |
| Age<sup>2</sup> | 5.90e-05 | 2.52e-05 | 24 | 2.34 | 0.028 |
| Hearing | 0.08 | 0.02 | 24 | 5.00 | < 0.001 |
| Speech material:Masker | -0.05 | 8.07e-03 | 328 | -6.33 | < 0.001 |
| SU:Age | 9.04e-06 | 4.11e-06 | 328 | 2.20 | 0.028 |
| SU:Speech material | -4.78e-04 | 1.57e-04 | 328 | -3.04 | 0.003 |
| Age:Masker | -5.68e-04 | 2.15e-04 | 328 | -2.64 | 0.009 |
| Hearing:Masker | -0.03 | 7.89e-03 | 328 | -3.48 | 0.001 |

In addition to this, we analyzed whether speech understanding or SNR best explains the differences in neural envelope tracking. We chose to compare two separate LMMs because evaluating whether adding SNR to the SU model resulted in an improved model fit could introduce problems with multicollinearity. A correlation analysis per speech material, masker and for the NH and HI group separately, revealed that 4 of the 8 Spearman correlations between the predictors SU and SNR had values higher than 0.70 (e.g. Matrix SWN (NH) = 0.77). Hence, we compared two non-nested LMMs with five predictors (SU or SNR, speech material, age, age^2^, hearing) and their interactions using AIC and Pseudo-R^2^ (e.g. SU model: envelope tracking ~ SU + speech material + age + age^2^ + hearing + SU:speech material + SU:age). The predictor “masker” was not included in these models because it would remove a large part of the variability that SNR could explain compared to SU. More specifically, in this study we used the SRT to create subject-specific SNRs. As can be inferred from figure 3, large differences in SNR were obtained for SWN and CT condition and thus including masker as a predictor in the SNR model, would not allow us to investigate the unique effect of SU versus SNR on envelope tracking. The comparison between the two LMMs revealed a smaller AIC and a higher Pseudo-R^2^ for the model in which SU was included as a predictor (AIC = −1425.57; Pseudo-R^2^ = 0.7086) compared with the model using SNR as a predictor (AIC = −1344.79; Pseudo-R^2^ = 0.6511), suggesting that envelope tracking is related to changes in speech understanding that cannot be explained by changes in SNR.

### 3.3 Neural consequences of hearing impairment

#### 3.3.1 Neural envelope tracking

In the subject-specific SNR conditions, HI adults demonstrated a significantly higher envelope tracking compared to their age-matched NH peers (figure 4; table 3), with a smaller difference for CT (mean difference between NH CT vs NH SWN = −0.0507, p = 0.007) than SWN (mean difference between NH SWN vs HI SWN = −0.0781, p < 0.001). Similarly for the fixed SNR conditions, statistics revealed a significantly increased envelope tracking for HI adults compared to their NH peers (p = 0.018; table 4). Taken together, these results demonstrate a significant enhanced neural envelope tracking of the target speech for adults with a hearing impairment, most apparent when speech was embedded in a stationary noise. To assess whether differences in SNR between NH and HI adults explain the enhanced effect of hearing impairment, we evaluated the LMM in which SNR was included as a predictor (see section above). Statistics revealed a significant main effect of hearing (p < 0.05) despite the fact that SNR was included in the model. This suggests that other factors than SNR also underlie enhanced envelope tracking in HI adults.

**Table 4:** Linear Mixed-effect Model: The effect of hearing impairment (hearing), fixed SNR condition (No noise versus 0 dB SNR), type of speech material and age on neural envelope tracking. Regression coefficients (*β* values), standard errors (SE), degrees of freedom (df), t-Ratios and p-values are reported per fixed-effect term. The variable participant nested in SU-level and speech material, was included as a random effect. Interactions are indicated by the symbol “:”.

| Fixed-effect terms | $\beta$ value | SE | df | t-Ratio | p-value |
| --- | --- | --- | --- | --- | --- |
| Intercept(for NH/Matrix/No noise) | 0.22 | 0.07 | 52 | 3.39 | 0.001 |
| condition | -0.06 | 0.01 | 25 | -5.73 | < 0.001 |
| Speech material | 0.04 | 0.03 | 52 | 1.16 | 0.253 |
| Age | -5.48e-03 | 2.74e-03 | 24 | -2.00 | 0.057 |
| Age<sup>2</sup> | 7.20e-05 | 2.70e-05 | 24 | 2.67 | 0.013 |
| Hearing | 0.04 | 0.02 | 24 | 2.53 | 0.018 |
| Speech material:Age | -1.31e-03 | 5.10e-04 | 52 | -2.56 | 0.013 |

#### 3.3.2 Target versus competing talker

In the CT condition, we also analyzed neural envelope tracking to the unattended, competing talker for the subject-specific SNR conditions (figure 5). This way, we could investigate if hearing impairment results in a differential neural tracking of target versus competing talker which can be considered a proxy for the ability to segregate talkers. We built a new LMM with five predictors (SU, speech material, age, hearing and attention (target versus competing); table 5). Post hoc analysis on the significant interaction between type of hearing and attention (table 5) indicated that both NH and HI adults obtained a higher neural envelope tracking for the target versus competing talker, with a larger neural segregation between talkers for the HI participants (mean difference for HI target vs competing = 0.0857, p < 0.001) compared to their NH peers (mean difference for NH target vs competing = 0.0364, p = 0.02). This differential neural envelope tracking is mainly due to the enhanced tracking of the target talker in HI adults because post hoc analysis revealed a significantly lower envelope tracking for the target talker for NH versus HI participants (mean difference = −0.0537, p = 0.005) and a non-significant, small difference between NH and HI adults for the competing talker (mean difference = −0.00446, p = 0.77).

**Figure 5:**
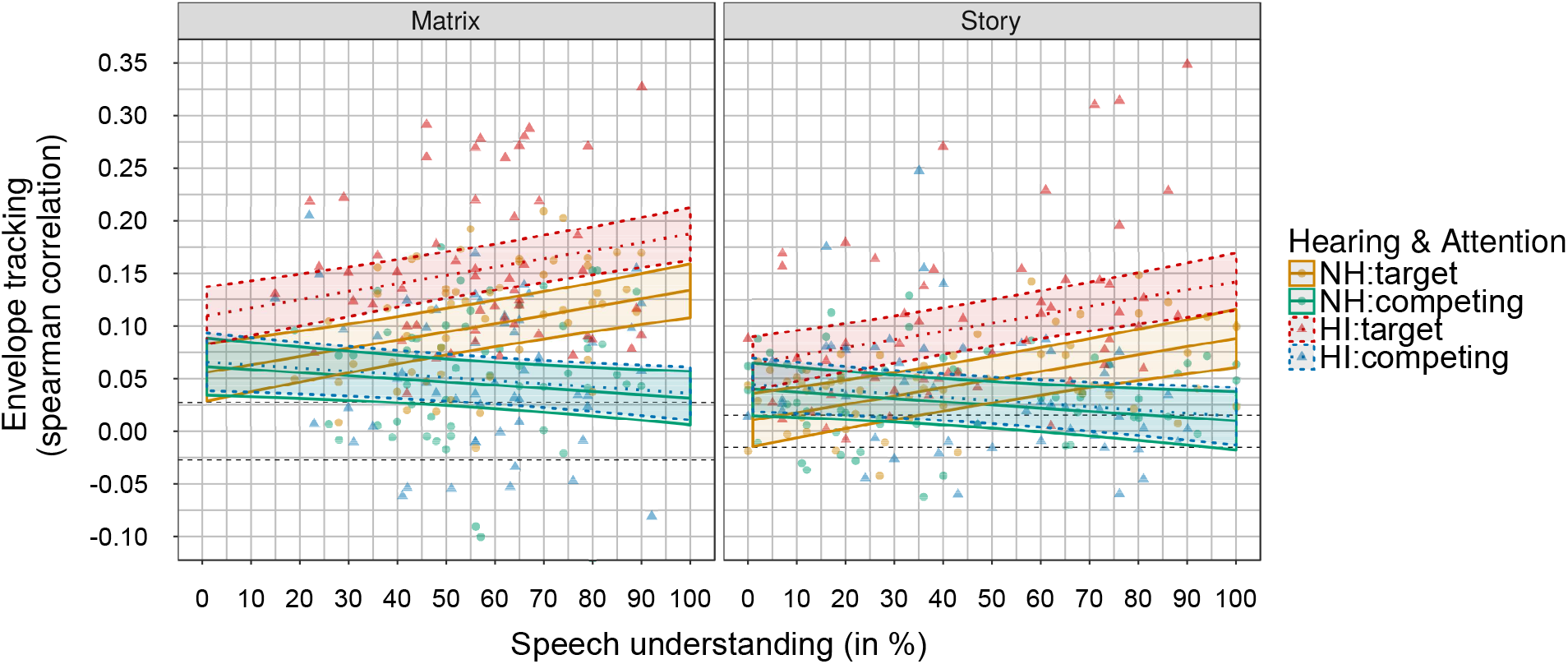
Neural envelope tracking of the target and competing talker as a function of speech understanding, for the matrix sentences and story (target) presented in competing speech (competing). Per speech material, a regression line was plotted on all subject-specific SNR data points for both hearing impaired (HI) and normal-hearing (NH) adults (color-coded). The dashed black lines indicate the significance level.

**Table 5:**
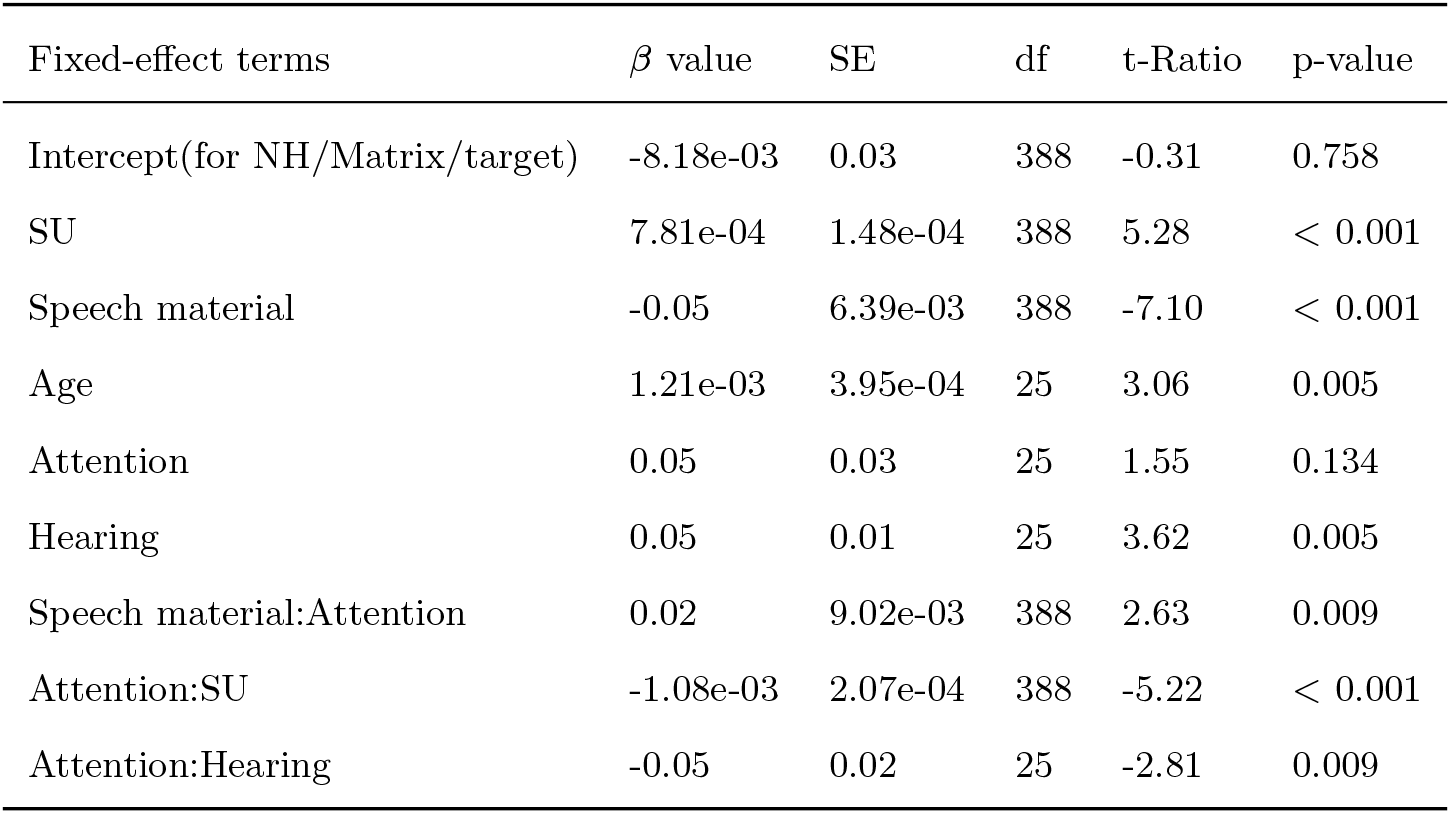
Linear Mixed-effect Model: The effect of speech understanding (SU), hearing impairment (hearing), attention (target versus competing), type of speech material (matrix vs story) and age on neural envelope tracking. Regression coefficients (*β* values), standard errors (SE), degrees of freedom (df), t-Ratios and p-values are reported per fixed-effect term. The variable participant nested in talker, was included as a random effect. Interactions are indicated by the symbol “:”.

| Fixed-effect terms | $\beta$ value | SE | df | t-Ratio | p-value |
| --- | --- | --- | --- | --- | --- |
| Intercept(for NH/Matrix/target) | -8.18e-03 | 0.03 | 388 | -0.31 | 0.758 |
| SU | 7.81e-04 | 1.48e-04 | 388 | 5.28 | < 0.001 |
| Speech material | -0.05 | 6.39e-03 | 388 | -7.10 | < 0.001 |
| Age | 1.21e-03 | 3.95e-04 | 25 | 3.06 | 0.005 |
| Attention | 0.05 | 0.03 | 25 | 1.55 | 0.134 |
| Hearing | 0.05 | 0.01 | 25 | 3.62 | 0.005 |
| Speech material:Attention | 0.02 | 9.02e-03 | 388 | 2.63 | 0.009 |
| Attention:SU | -1.08e-03 | 2.07e-04 | 388 | -5.22 | < 0.001 |
| Attention:Hearing | -0.05 | 0.02 | 25 | -2.81 | 0.009 |

In addition, we found an increase in envelope tracking of the target talker with increasing speech understanding (table 5; table 3) but no significant change of the competing talker (p > 0.05; table 5). Furthermore, the difference in tracking between target and competing talker was larger for the matrix sentences (mean difference for matrix target vs competing = 0.0729, p < 0.001) than the story (mean difference for story target vs competing = 0.0492, p < 0.001). Finally, a significant main effect of age was found but no further interactions, suggesting an increase in neural envelope tracking for both target and competing talker with advancing age (p = 0.005; table 5).

#### 3.3.3 Cortical ASSRs to tone pips

In addition to measuring neural tracking of speech stimuli, we also evaluated whether hearing impairment influenced the neural SNR of cortical ASSRs to tone pips (figure 6). The Linear Fixed-effect Model (LFM) did not reveal a significant effect of hearing impairment (*β* = 0.81, SE = 1.14, p = 0.480) or age (*β* = −0.06, SE = 0.03, p = 0.070) on neural envelope tracking.

**Figure 6:**
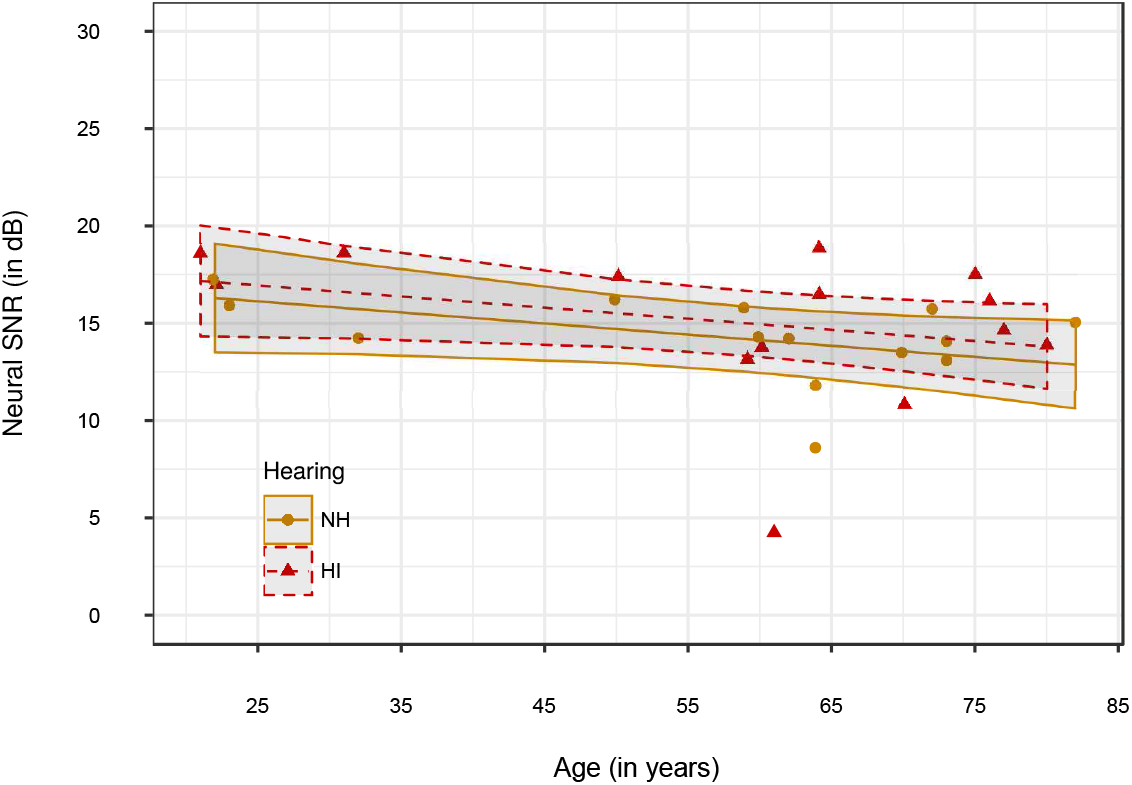
The neural SNR of ASSRs to tone pips as a function of age. Regression lines with confidence intervals (shaded areas) were fitted on all the data points, for both normal-hearing (NH) and hearing impaired (HI) adults (color-coded).

### 3.4 The interplay between neural envelope tracking, cognition and self-reported listening effort

Research has demonstrated that hearing loss can be associated with a worse performance on cognitive tasks, even without having a cognitive impairment. To investigate if the enhanced neural envelope tracking of the target talker for HI versus NH adults is related with cognitive deficits, we compared the results of NH and HI adults on two cognitive tests, the Reading Span test (RST) and Stroop test, using a LFM with two predictors, age and hearing, per test. Similar to our previous study (Decruy et al., 2019), we found a significant decrease in scores with advancing age for both the RST (*β* = −0.26, SE = 0.06, p < 0.001) and Stroop test (*β* = −0.74, SE = 0.34, p = 0.040), but no significant effect of hearing impairment (RST: p = 0.100; Stroop: p = 0.260).

Next to cognition, research has shown that in general HI adults need to spend more listening effort to understand speech in challenging situations compared to their NH peers (Ohlenforst et al., 2017). To assess if the enhanced envelope tracking for HI adults reflects increased listening effort, we analyzed the listening effort ratings that participants reported for each SNR condition during the EEG experiment. For the subject-specific SNR conditions, we did not find a significant effect of hearing impairment (LMM with only hearing as predictor: p = 0.906). For the fixed SNR conditions, on the other hand, we found that HI adults reported significantly more effort compared to their NH peers (*β* = 24.15, SE = 6.36, p = 0.001). In spite of this, likelihood ratio tests and AIC did not show a significant contribution of effort when adding the self-reported effort scores, as a main or interaction effect, to the best fitting model for envelope tracking at the fixed SNRs.

## 4. Discussion

We investigated the effect of hearing impairment on neural envelope tracking by comparing the results of 14 HI with 14 age-matched NH adults. Both NH and HI adults showed an increase in envelope tracking of the target talker with increasing speech understanding. Interestingly, our results suggest that HI adults can neurally segregate the target and competing talker, but need an additional enhancement of the target talker compared to their NH peers, to obtain this. Lastly, no significant effect of hearing impairment was detected on cortical ASSRs.

### 4.1 Hearing impairment is more detrimental when speech is embedded in a competing talker

Prior to measuring neural envelope tracking, we also assessed the effect of hearing impairment on speech understanding in noise. In line with previous studies, our behavioral results demonstrate that persons with sensorineural hearing loss obtain less benefit from listening in the gaps of a fluctuating masker, despite having access to amplification (Festen and Plomp, 1990; Goossens et al., 2017; Shinn-Cunningham and Best, 2008). In other words, sensorineural hearing loss results in temporal processing deficits which can not be overcome by simply amplifying the signal. Degradation due to stationary noises, on the other hand, seems to be more easy to compensate using simple linear amplification, as we found smaller differences between the matrix SRTs of NH versus HI adults.

### 4.2 Hearing impaired adults show enhancement of the target talker when neurally segregating speech from noise

#### 4.2.1 Comparison with other neural envelope tracking studies

For both NH and HI adults, neural envelope tracking was measured to sentences and a story, both presented in stationary speech-weighted noise and a competing talker. In line with the three previous studies, HI listeners showed higher envelope tracking to the target than the competing talker (Mirkovic et al., 2019; Petersen et al., 2017; Presacco et al., 2019), indicating that HI adults can neurally segregate different talkers despite having a disabling hearing loss. However, we observed a different effect of hearing impairment on neural envelope tracking of the target or competing talker than these previous studies.

Petersen et al. (2017) measured neural envelope tracking to stories in HI adults with varying degrees of hearing loss. Their results show that hearing loss resulted in an enhanced tracking of the competing talker but no alteration of the target talker. In other words, severe hearing impariment resulted in a weak differential tracking of target versus competing talker. In the present study, however, HI participants showed, compared to their age-matched NH adults, a significantly enhanced envelope tracking of the target talker when speech was embedded in a stationary noise or competing talker. Since no enhancement was found for the competing talker, our results suggest, in contrast to Petersen et al. (2017), that HI adults require a larger differential tracking of target versus competing talker to neurally segregate speech from noise. We hypothesize that this disagreement can be due to methodological differences. We analyzed the effect of hearing impairment on the raw correlations between the reconstructed and actual envelope whereas Petersen et al. (2017) performed several post-processing steps on their EEG-data, limiting their analysis to clusters of EEG channels and time windows (time lags corresponding with CAEPs components). In addition, we controlled for the effects of age by carefully age-matching our NH and HI participants while Petersen et al. (2017) eliminated the effects of age on their PTA measure but not on neural envelope tracking, which is also known to be affected by age (Decruy et al., 2019; Presacco et al., 2016). Furthermore, the *quasi*-linear hearing aid algorithm could have affected the envelope in a different way than our controlled linear amplification (see 2.2.1).

Next to Petersen et al. (2017), our results are not entirely in line with the findings of Presacco et al. (2019). They measured neural envelope tracking in NH and HI older adults and found no effect of hearing impairment on the tracking of nor the target nor competing talker. A plausible explanation for the difference for the target talker could result from the fact that Presacco et al. (2019) could not carefully age-match their participants. The HI adults were significantly older than their NH adults which could result in a more dominant effect of age over hearing impairment. In addition to age-matching, Presacco et al. (2019) measured neural envelope tracking using MEG instead of EEG responses. Taking into account that MEG is mainly dominated by cortical sources whereas EEG is sensitive to both subcortical and cortical regions (Lopes da Silva, 2013), enhanced envelope tracking in our HI adults might reflect enhanced brainstem responses. Although this may seem a plausible hypothesis, we have to note that our EEG responses were filtered between 1 and 8 Hz and thus should mainly reflect cortical responses. Nevertheless, it seems useful to record brainstem responses to natural speech (Etard et al., 2019) or perform source or connectivity analyses because recent studies have suggested that hearing impairment alters the relation between subcortical and cortical levels of the auditory system (Bidelman et al., 2014; Presacco et al., 2019). Another reason for the difference between our results and Presacco et al. (2019), could involve the fact that they did not use subject-specific amplification. Lastly, Mirkovic et al. (2019) also measured neural envelope tracking in NH and HI adults. In line with our results, no effect of hearing impairment was revealed on the tracking of the competing talker. With regard to the target talker, no statistics were reported, but their results suggest a higher envelope tracking for the target talker for HI adults compared to NH counterparts (figure 7 in Mirkovic et al., 2019).

#### 4.2.2 Possible explanations underlying enhanced envelope tracking in hearing impaired adults

Several mechanisms could underlie the enhanced envelope tracking to the target talker in our HI adults. A first plausible explanation involves the reported enhanced envelope sensitivity in persons with a hearing loss. Several behavioral studies have demonstrated lower, and thus better, amplitude detection thresholds for HI versus NH adults (Füllgrabe et al., 2003; Wallaert et al., 2017) as well as an enhanced sensitivity to suprathreshold modulations (Moore et al., 1996). Accordingly, Millman et al. (2017) showed enhanced envelope encoding in HI adults compared to their NH peers when measuring MEG responses to amplitude modulated noise in humans with a sensorineural hearing loss. Although it is plausible that the increased envelope tracking in HI reflects enhanced envelope sensitivity, we cannot entirely confirm this hypothesis since we did not link the observed neural enhancement in our study to modulation detection thresholds or suprathreshold modulation sensitivity in the same participants. Future research is thus needed to verify this.

Since we presented our speech stimuli at higher SNRs for HI adults to achieve a wide range of speech understanding levels per individual, differences in SNR might be a second explanation for driving the differences in envelope tracking between NH and HI adults. However, we do not think that this is likely because of the following reasons. Firstly, our results reveal a larger enhancement in neural envelope tracking when speech was presented in SWN, where more similar SRTs were obtained between NH and HI adults. Secondly, our results also demonstrate significantly enhanced envelope tracking of the target talker for the fixed SNR conditions (No noise, 0 dB SNR), ruling out the possibility that differences in SNR would explain the neural effect of hearing impairment. Lastly, we found that the predictor speech understanding explains more variability in envelope tracking than SNR and that the effect of hearing loss remains when including SNR instead of SU as a predictor in the LMM.

A last possible explanation is based on a cognitive perspective. Previous research has indicated that older persons without cognitive impairment but disabling hearing loss, perform worse on cognitive tests (Humes et al., 2013; Lin et al., 2011). In addition, studies have also shown that HI adults need more effort to perform in the same way as their NH counterparts (Ohlenforst et al., 2017). Hence, the additional enhancement associated with hearing impairment could reflect a compensation of HI adults to separate target speech from background noise. To investigate this, we assessed whether differences between NH and HI adults for cognitive results and effort ratings would explain the observed enhanced envelope tracking.

Similar to Presacco et al. (2019), no significant relationship with cognitive results or effort ratings was found in the present study. This may be due to the possible lack of sensitivity of the measures that we have used. With regard to cognition, we have used a visual-orientated Stroop and Reading Span test whereas studies have suggested that auditory-cognitive tests, such as a Listening span test, reveal differences more easily (e.g. Smith et al., 2016). However, we have to note that these researchers could not entirely rule out differences in age between the NH and HI group. With regard to listening effort, there is a great debate about how to define and quantify effort. This is partly due to the large variety of measures that are being used (Alhanbali et al., 2019; McGarrigle et al., 2014; Pichora-Fuller et al., 2016). Although self-report measures are easy to administer and widely used, several studies have failed to find a robust link between self-reported effort and behavioral or physiological effort measures (Anderson Gosselin and Gagné, 2011, Zekveld et al. (2011); Decruy et al., n.d.; Gagné et al., 2017) as well as neural envelope tracking measures (Decruy et al., n.d.; Müller et al., 2019). To verify whether the enhanced envelope tracking in HI adults is related to listening effort, it would be beneficial to also include other effort measures, such as behavioral reaction times, alpha power or pupillometry.

In conclusion, our results do not demonstrate that envelope tracking can be used to quantify listening effort, i.e. how much brain resources are deliberately recruited to track the speech envelope. Nevertheless, we obtained different results for the target versus competing talker, indicating that our neural envelope tracking measure is a good measure of how individuals neurally segregate speech from background noise. In view of this, we do not completely rule out the cognitive, compensatory hypothesis because the observed, enhanced envelope tracking of the target talker could suggest that HI adults require a larger differential tracking of target versus competing talker in order to neurally segregate speech from background noise. As mentioned earlier, research has suggested that hearing impairment leads to a higher interdependence between subcortical and cortical levels of the auditory system (Bidelman et al., 2014; Presacco et al., 2019). Taking this into account, enhanced tracking of the target talker could reflect a neural mechanism that compensates for degraded responses at the subcortical level. To confirm this hypothesis, furture studies should be conducted in which source analyses and recordings of brainstem responses to natural speech are included.

#### 4.2.3 Differential effect of hearing impairment on cortical ASSRs

In contrast to neural envelope tracking, we did not find a significant effect of hearing impairment on cortical ASSRs. This was unexpected since Millman et al. (2017) showed enhanced MEG responses to modulated noise for adults with sensorineural hearing loss. This disagreement could be explained based on a recent study which suggests that differences in ASSRs between NH and HI adults change with advancing age. More specifically, Goossens et al. (2019) showed enhanced cortical and subcortical ASSRs in young (20-30 years) and middle-aged (50-60 years), but not in older HI adults (70-80 years). Although our NH and HI participants were carefully age-matched, we mainly included adults older than 55 years (20 out of 28). In addition, our sample size per group was relatively small (14 NH and 14 HI adults) which makes it more difficult to detect small effects. We believe that these reasons could also explain why our current results did not reveal a significant decrease in cortical ASSRs or consistent supralinear increase in envelope tracking with advancing age, as shown in our previous study (Decruy et al., 2019). Hence, increasing the sample size and measuring neural envelope tracking in more young and middle-aged HI adults will allow further disentanglement of the effects of age versus hearing impairment. We have to note, however, that this will not be straight forward as the prevalence of sensorineural hearing loss in young and middle-aged adults is low.

Another explanation stems from the findings of neuro-imaging studies which demonstrate a cortical reorganization for persons with a hearing impairment. For example, Du et al. (2016) showed that adults with a mild hearing loss show increased activity in prefrontal cortices during a speech identification task. Similarly, Campbell and Sharma (2013) showed that HI adults show increased amplitudes for CAEPs for frontal brain regions whereas temporal regions showed decreased neural activity compared to NH peers. Taking this into account, different brain regions may be involved in evoking cortical ASSRs to tone pips versus neural envelope tracking in HI adults. Therefore, it would be beneficial to perform a source analysis to get more insight into which brain regions evoke enhanced neural envelope tracking in HI adults.

Lastly, we hypothesize that differences in the characteristics of stimuli can cause different effects of hearing impairment. In the present study, we presented tone pips. In other studies noise has been used that is amplitude modulated by different carrier signals (sine modulator: Goossens et al., 2019; square-wave modulator: Millman et al., 2017) to elicit ASSRs. Furthermore, we hypothesize based on the results of Koerner and Zhang (2018) that differences in the processing of simple tone pips versus speech can also underlie the differential effect of hearing impairment on neural envelope tracking versus cortical ASSRs. Koerner and Zhang (2018) measured speech-evoked CAEPs and mismatch negativity (MNM) in 18 listeners with and without hearing loss. HI adults showed differences in the MNM but not the CAEPs compared to NH adults, suggesting that hearing impairment mainly affects later stages of auditory processing. As understanding speech involves much more high-level later processing compared to the processing of tone pips (e.g. mapping from acoustics to words), this may explain why we did not observe a significant difference between NH and HI adults for cortical ASSRs.

### 4.3 Neural envelope tracking increases with speech understanding

For both speech materials and maskers, we found an increase in envelope tracking of the target talker with increasing speech understanding for NH, but also HI adults. Put otherwise, our results demonstrate that an individual with a hearing impairment shows higher envelope tracking when speech is more intelligible, despite having an overall enhanced envelope tracking. This is not entirely in agreement with Petersen et al. (2017) who found that hearing loss impairs the association between neural envelope tracking and changes in SNR. However, they only compared the most favorable condition (No noise) with the most difficult condition (SRT for 80% SU - 4 dB) and did not take into account the potential distortion of the original envelope by the hearing aid. Presacco et al. (2019), on the other hand, found an increase in envelope tracking as a function of SNR for HI adults, but not for older NH adults. The latter can be due to the fact that all presented SNRs were highly intelligible for the NH adults. In sum, our findings provide further support for the value of neural envelope tracking to objectively measure speech understanding (Decruy et al., 2019; Lesenfants et al., 2019; Vanthornhout et al., 2018).

Next to diagnostics, measuring neural envelope tracking could also be useful to evaluate the benefit of hearing aids. Recently, Karawani et al. (2018) have shown that new hearing aid users already show altered cortical responses to a speech syllable, within six months of hearing aid use. As daily life communication involves mainly natural speech, future studies should investigate how neural envelope tracking evolves over time in hearing aid users. We have to note, however, that current hearing aids not only amplify but also compress the incoming sound. Since we provided linear frequency-specific amplification, there is a need for future research that assesses the link between speech understanding and neural envelope tracking to the compressed speech envelope.

## 5. Conclusion

The present study demonstrated that hearing impaired adults can achieve a robust neural segregation of speech versus background noise. Furthermore, our results demonstrate an enhancement of envelope tracking of the target talker for persons with sensorineural hearing loss compared to age-matched normal-hearing adults. This could suggest that hearing impaired adults have an enhanced sensitivity to envelope modulations or need a larger difference between speech and background noise in order to neurally segregate different streams. Finally, we provided further support for neural envelope tracking to objectively measure speech understanding as our HI adults also showed an increase in envelope tracking when understanding speech better. This could be particularly useful towards the applications of neuro-steered hearing aids as well as the objective evaluation of speech understanding performance in difficult-to-test populations, such as hearing impaired children.

## Abbreviations

AIC: Akaike’s Information Criterion
ASSR: Auditory steady-state response
CAEP: Cortical auditory evoked potentials
CT: competing talker
dB HL: decibel Hearing level
dB SNR: decibel Signal-to-noise ratio
df: degrees of freedom
HI: hearing impairment
EEG: electroencephalography
LFM: Linear Fixed-effect Model
LMM: Linear Mixed-effect Model
MEG: magnetoencephalography
NAL-RP: National Acoustics Laboratory-Revised Profound algorithm
NH: normal-hearing
PTA: pure tone average
RST: Reading Span Test
SE: standard error
SNR: signal-to-noise ratio
SRT: speech reception threshold
SU: speech understanding
SWN: speech-weighted noise1

## Acknowledgments

Our special thanks goes to all participants for their cooperation in this study. We also want to ackownledge the help of master’s students Elien van den Borre and Mélissa Schoubben in participant recruitment and data acquisition. Furthermore, we want to thank audiologists Mieke Goossens, Lise Goris and Andre Bussé for their cooperation in the recruitment of the hearing impaired adults. Lastly, we also want to acknowledge the section editor and two anonymous reviewers for their help in improving the manuscript.

